# 3D imaging of colorectal cancer organoids identifies responses to Tankyrase inhibitors

**DOI:** 10.1101/705277

**Authors:** Luned M. Badder, Andrew J. Hollins, Bram Herpers, Kuan Yan, Kenneth B. Ewan, Jennifer R. Shone, Delyth A. Badder, Marc Naven, Kevin Ashelford, Rachel Hargest, Alan R. Clarke, Christina Esdar, Hans-Peter Buchstaller, Dirk Wienke, Leo S. Price, Paul Shaw, Trevor C. Dale

**Affiliations:** Cardiff University School of Biosciences, Cardiff, Wales, U.K.; European Cancer Stem Cell Research Institute (ECSCRI), Cardiff University, Wales, U.K.; OcellO B.V Leiden, The Netherlands; Cellular Pathology Department, University Hospital for Wales, Cardiff, UK; Division of Cancer and Genetics, School of Medicine, Cardiff University, Cardiff, UK; Department of Colorectal Surgery, University Hospital of Wales, Cardiff, UK; CCMRC, Division of Cancer and Genetics, Henry Wellcome Building, Cardiff University, Cardiff, UK; Merck Healthcare KGaA, Biopharma, Research & Development, Frankfurter Str. 250, 64293, Darmstadt, Germany; Velindre Cancer Centre, Cardiff, Wales, U.K.

**Keywords:** Organoid, Tankyrase inhibitors, High content imaging, Wnt, Colorectal cancer, cancer stem cells

## Abstract

Aberrant activation of the Wnt signalling pathway is required for tumour initiation and survival in the majority of colorectal cancers. The development of inhibitors of Wnt signalling has been the focus of multiple drug discovery programs targeting colorectal cancer and other malignancies associated with aberrant pathway activation. However, progression of new clinical entities targeting the Wnt pathway has been slow. One challenge lies with the limited predictive power of 2D cancer cell lines because they fail to fully recapitulate intratumoural phenotypic heterogeneity. In particular, the relationship between 2D cancer cell biology and cancer stem cell function is poorly understood. By contrast, 3D tumour organoids provide a platform in which complex cell-cell interactions can be studied. However, complex 3D models provide a challenging platform for the quantitative analysis of drug responses of therapies that have differential effects on tumour cell subpopulations. Here, we generated tumour organoids from colorectal cancer patients and tested their responses to inhibitors of Tankyrase (TNKSi) which are known to modulate Wnt signalling. Using compounds with 3 orders of magnitude difference in cellular mechanistic potency together with image-based assays, we demonstrate that morphometric analyses can capture subtle alterations in organoid responses to Wnt inhibitors that are consistent with activity against a cancer stem cell subpopulation. Overall our study highlights the value of phenotypic readouts as a quantitative method to asses drug-induced effects in a relevant preclinical model.

## Introduction

Aberrant canonical Wnt/β-catenin signalling, as a result of activating mutations within the pathway, has a prominent role in the initiation and progression of colorectal cancer (CRC) [1, 2]. A number of feedback loops control β-catenin turnover and Wnt activation. In the absence of a Wnt ligand the multi-protein β-catenin destruction complex, formed of AXIN1/2, Adenomatous polyposis coli (APC) and glycogen synthase kinase (GSK3β), mark β-catenin for degradation [3]. As a result, the accumulation and subsequent translocation of β-catenin to the nucleus is inhibited, preventing the downstream activation of target genes [4]. Components of the destruction complex are also tightly regulated. AXIN1 and AXIN2 are concentration-limiting components of the destruction complex. Levels of AXIN1/2 are post-transcriptionally regulated by tankyrases (TNKS1 and TNKS2), members of the poly (ADP-ribose) polymerases (PARP) family of enzymes, which enhance Wnt signalling by targeting AXIN1/2 for degradation [5].

The inhibition of Wnt signalling has been validated as a means of blocking tumour growth in many cancer models [6, 7]. Small molecule inhibitors of TNKS1 and TNKS2 have been shown to reduce Wnt signalling in intestinal cancer cell lines and have been suggested to prevent tumour growth due to their ability to stabilise AXIN1 and AXIN2 levels and as a result, to inhibit β-catenin mediated transcription [5, 8–16]. However, the progression of TNKSi into clinical trials has been restricted due to significant issues of intestinal toxicity within *in vivo* models, emphasising the important role of Wnt signalling in adult tissue homeostasis [17] [18]. Furthermore, the TNKSi described above reduced colorectal cancer cell numbers in 2D culture, but did so with relatively low effect sizes by comparison with their ability to reduce levels of Wnt/TCF-dependent transcription in the same cell lines. Similar partial-efficacy cell line responses were observed during the development of inhibitors of the CDK8 and CDK19 kinases, suggesting that growth in 2D cell culture may not be an optimal readout for compounds that are anticipated to target cancer stem cells [7].

To date, most preclinical *in vitro* studies have relied on the use of conventional 2D cultures of cell lines. Whilst cancer cell lines can be usefully used to study pathway deregulation, they frequently fail to be predictive when used as readouts of antitumour efficacy [19]. It is well recognised that immortalised 2D cell lines adapted to growth on plastic poorly represent the intricate cellular cross-talk present in tumours. In particular, they are unable to recapitulate the inter-cellular interactions that form cancer stem cell niches and, as a consequence, such models fail to fully reflect *in vivo* results.

3D tumour organoids have been shown to provide a more complex insight into tumour cell-cell interactions [20, 21] and have been recognised as models with the potential to bridge the gap between *in vitro* and *in vivo* preclinical studies. Organoids that have never been adapted for growth on plastic have been cultured from multiple tumour types, and have been shown to better represent the genetic diversity of distinct tumour subtypes than 2D cell lines [22]. Organoids derived from genetically-engineered mice in which the Wnt pathway had been oncogenically-activated accurately predicted subsequent *in vivo* responses to inhibitors of the CDK8 and CDK19 kinases, while 2D cell culture only showed partial-efficacy [7]. Organoids have been further shown to predict patient responses and might in future be used in the clinic in personalised medicine [23].

The biological and phenotypic complexity of 3D primary organoid cultures allow, in principle, the assessment of compound activity against a subset of inter-cellular signalling that is central to tumour growth. However, most uses of organoid assays to date have relied on fixed end-point metabolic assays (e.g. ATP-level quantification) that aggregates responses in every cell within a population of organoids. To fully exploit the potential of organoid assays, analysis platforms need to yield quantitative data that can clearly represent compound-induced effects on signalling pathways whose outputs are complex, whilst simultaneously maintaining usability in a high throughput format.

In this work, we generated CRC patient-derived organoids and studied their responses to Tankyrase inhibitors (TNKSi). Using metabolic end point assays, tankyrase inhibition showed partial efficacy, reflecting a limited reduction in the growth of TNKSi-sensitive organoids. Closer examination of organoid responses suggested that TNKSi altered the ratio of stem-like to differentiated cell populations. This subtle effect on the cellular composition of organoids was best identified by multiparametric imaging analysis of organoids that was nonetheless compatible with high throughput analysis. Our findings demonstrate the potential of a phenotypic approach to assess drug-induced phenotypes in a preclinical setting, which may be applicable to a wider range of therapeutics that target cancer stem cell biology and niches.

## Results

### Establishment of Tumour organoids from CRC patient material

Surgically-resected CRC material was isolated from patients under informed consent. Cell fragments were isolated from tissue and plated within growth factor-reduced matrigel within 24 hours following surgery, adapting methods previously described by Sato et al [20]. Two different culture conditions were used as standard; “7+ basal” and “full” growth factor rich medium. Each condition was found to be optimal for a distinct subset of organoids (Figure 1A) and allowed the efficient generation of organoids from what were in some cases limited tumour cell numbers. Of 59 collected samples 48 were established in culture, a ‘take-rate’ of >81% for organoid model derivation (organoids were denoted by Isolation number, ‘Iso-’). Recent studies have similarly emphasised the requirement for different media to ensure efficient organoid generation from different patients [21].

**Figure 1.**
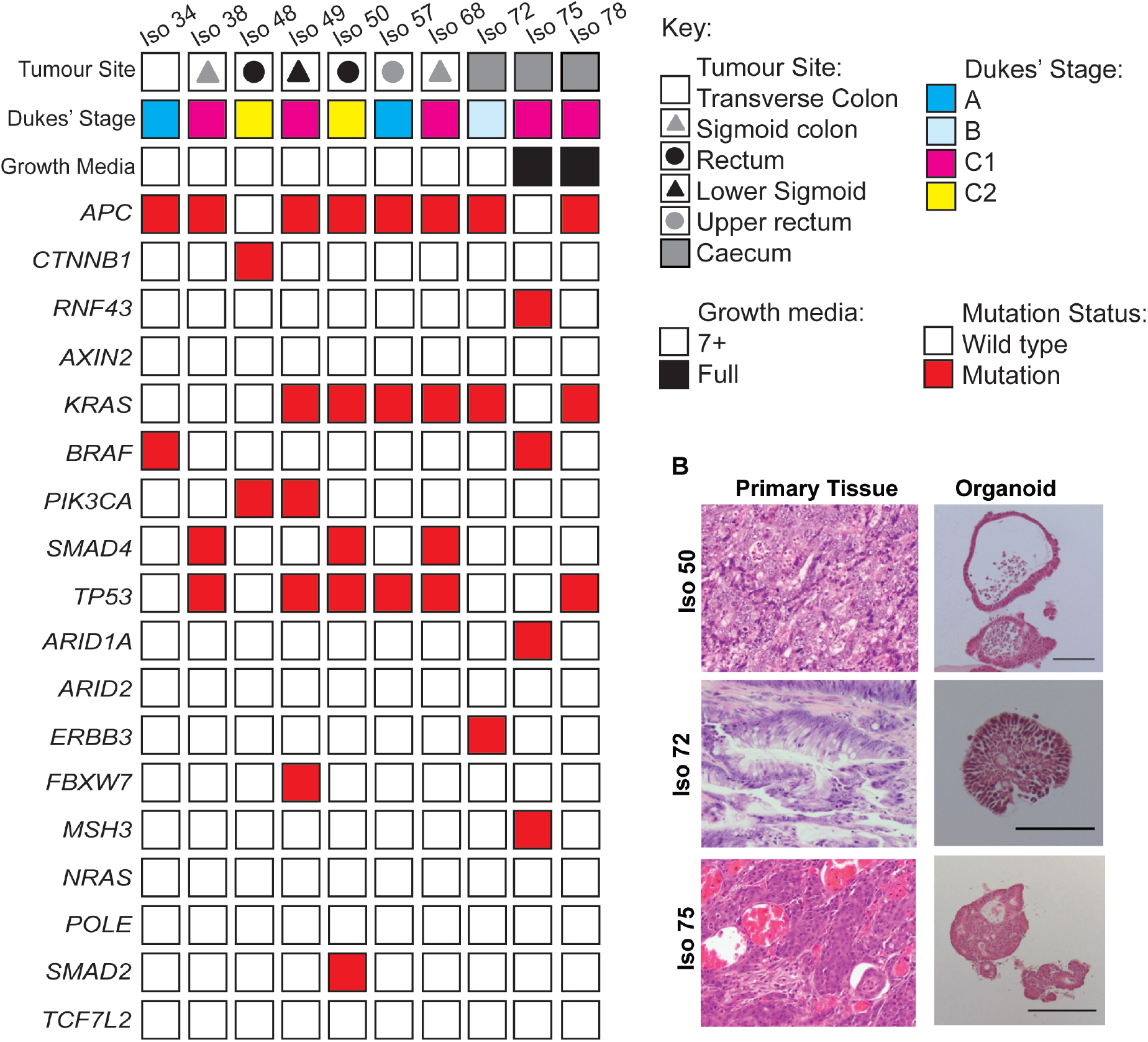
Generation and establishment of tumour organoids from CRC patient material. **A.** Summary of parental tumour sites and Dukes’ Stages, organoid media conditions, and mutation status of organoid lines identified in Whole Exome Sequencing analysis. **B.** Representative Hematoxylin+Eosin (H&E) staining of FFPE tissue sections from primary colorectal tumour patient material and organoid counterparts. Organoids were cultured for a minimum of 6 days prior to fixation and embedding. Qualitative morphological similarities were observed between tumour and organoid pairs. Scale bar = 100 μm.

A subset of 10 organoids was studied in more detail. Optimal culture conditions were determined on the basis of organoid formation efficiencies and visual inspection of organoid integrity. 8/10 organoid lines showed good growth in 7+ basal medium whilst 2/10 were dependent on the additional inclusion of EGF (50 ng/ml), Noggin (100 ng/ml), Nicotinamide (10 mM), A-83-01 (500 nM), SB202190 (10 μM), as well as Wnt3a- (40%) and R-spondin 1- (10%) conditioned media. The organoid: medium combinations shown in Figure 1A were used for subsequent studies. Full exome sequencing showed that the subset of 10 organoids contained common genotypes found in other organoid banks (Figure 1A, Supplementary table S1); [2, 24]. In particular, 80% of organoid lines harboured mutations in components of the Wnt signalling pathway, such as *APC* and β -*catenin (CTNNB1)*. Iso 75 was found to harbour a mutation in *RNF43*, which is associated with an increased Frizzled (Fz) receptor expression and Wnt ligand dependence [24, 25].

CRC organoids have previously been shown to display histological features in common with the tumours from which they were derived. Paired histological samples from the organoids described were compared to their parental tumours. Distinct morphologies were observed ranging from lines that preferentially formed single epithelial cell layers with large lumens containing apoptotic cells (Figure 1B, Supplementary Figure S1) to multi-layered organoids with small indistinct lumens. The organoids frequently retained key histopathological grading features that were present within the original patient tumour epithelium. For example, Iso 72, was established from a mucinous adenocarcinoma and displayed a high percentage of mucinous vacuole structures that resembled that of the matching patient tissue. The retention of features present within primary patient tumour tissue showed that many features of tumour morphology could be retained without the need for a stromal compartment.

### Tankyrase inhibition altered the growth of patient-derived organoids

To study CRC organoid responses to a Wnt pathway inhibitor, we utilised the tankyrase inhibitor (TNKSi), C1 (Figure 2A). After six days of treatment, AXIN2 and TNKS1/2 protein levels were raised in a number of organoid lines (Figure 2B). TNKS normally targets AXIN2 and TNKS1/2 for degradation and TNKSi treatment would be expected to stabilise levels of each protein [5]. To assess the functional impact of Tankyrase inhibition on organoid growth, cell viability assays were performed for 3 organoid lines; Iso 50, Iso 72 and Iso 75. Organoids were digested to near-single cells, embedded within growth-factor reduced Matrigel and overlaid with growth media containing the TNKSi for six days prior to measuring relative ATP levels (Cell Titer Glo 3D). C1, which has a cellular mechanistic EC_50_ of 2nM, showed partial efficacy in reducing ATP levels and markedly different potency in the 3 lines (Figure 2C-G). By comparison, the MEK inhibitor Trametinib induced strong apoptosis and necrosis and was associated with a robust efficacy assay window (data not shown). EC_50_ values ranged from 2nM to 1 μM, but accurate values were hard to establish due to assay variance and also because many cells survived, even in the presence of high concentrations of compound, leading to reduced assay windows. The partial survival of organoid cells after 6 days C1 treatment suggest that long-term AXIN2 and TNKS1/2 biomarker responses might be compromised by alterations in cellular composition.

**Figure 2.**
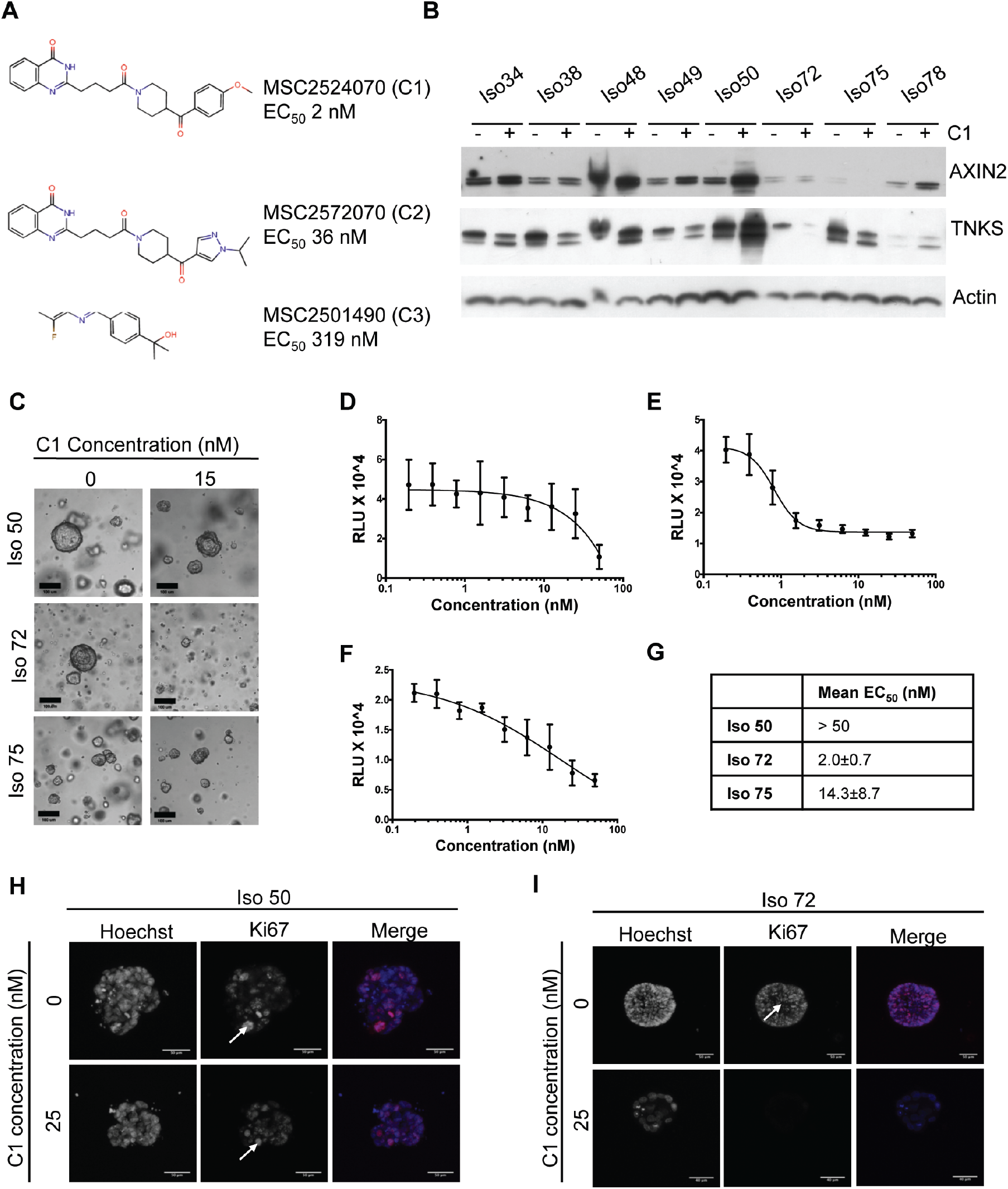
Assessing the effects of Tankyrase inhibition on overall organoid viability. **A.** Structures of three tankyrase inhibitors: MSC2524070A (C1), MSC2572070A (C2) and MSC2501490A (C3), with corresponding EC_50_ values. **B.** Western blot of eight organoid lines treated with C1 (15 nM) and harvested after 6 days. **C.** Representative images of organoids treated with C1 (15 nM) or DMSO (scale bar=100 μm). **D,E,F**. Dose-response curves of organoids treated with C1 for six days. Organoids previously established in suitable culture conditions were enzymatically digested to single cells and overlaid with media containing a dose range of drug (0.195 nM – 50 nM), or a DMSO control (0.1% in media). Cell Titer Glo 3D readouts were performed in triplicates for Iso 50 (**D**), Iso 72 (**E**), Iso 75 (**F**). **G.** EC_50_ values, obtained from (n=3) experiments. **H,I**. Representative immunofluorescent staining of a proliferation marker in organoids. Iso 50 (**H**) and Iso 72 (**I**) with differential sensitivity to TNKSi were treated with C1 (25 nM) or DMSO (0.1%) control for 6 days, fixed and stained with Ki67. Arrows indicate proliferating cell within the organoid, Ki67 positive-cells in Iso 72 only when treated with C1.

Immunofluorescent staining showed reduced levels of the proliferation marker Ki67 in Iso 72 but not Iso 50 organoids suggesting that reduced proliferation contributed to the lower ATP levels observed (Figure 2H,I).

### Multi-parametric phenotypic profiling of organoids in 3D to classify responses to Tankyrase inhibition

To better understand the TNKSi partial efficacy, a novel multi-parametric 3D image workflow analysis was used to characterise the morphological effects of the responses. Three Tankyrase 1/2 (TNKS1/2) small molecule inhibitors were used that differed over 3 orders of magnitude in their potency (C1; 2nM, C2; 36nM, C3; 319nM Figure 2A). Organoids were seeded and treated as previously for 6 days prior to fixation and staining with rhodamine-phalloidin (F-actin) and Hoechst 33258 (DNA). Images were acquired using a high-content wide-field fluorescence microscope as described previously [26, 27].

To characterise phenotypes, OMiner™ software (OcellO™) was used to extract morphological features from 2D projections of the F-actin and nuclei-derived image stacks to generate masks of individual organoids, internal lumen structures and nuclei. Image segmentation was used to distinguish components of both channels and minimise any background signal (Figure 3A, Supplementary Figure S4). Overall shape and fluorescence intensity of individual organoid structures were extracted on a well-by-well basis to generate approximately 600 quantified features. Such features included information in regards to shape, size, boundaries and branching of organoid lumens and nuclear shape.

**Figure 3.**
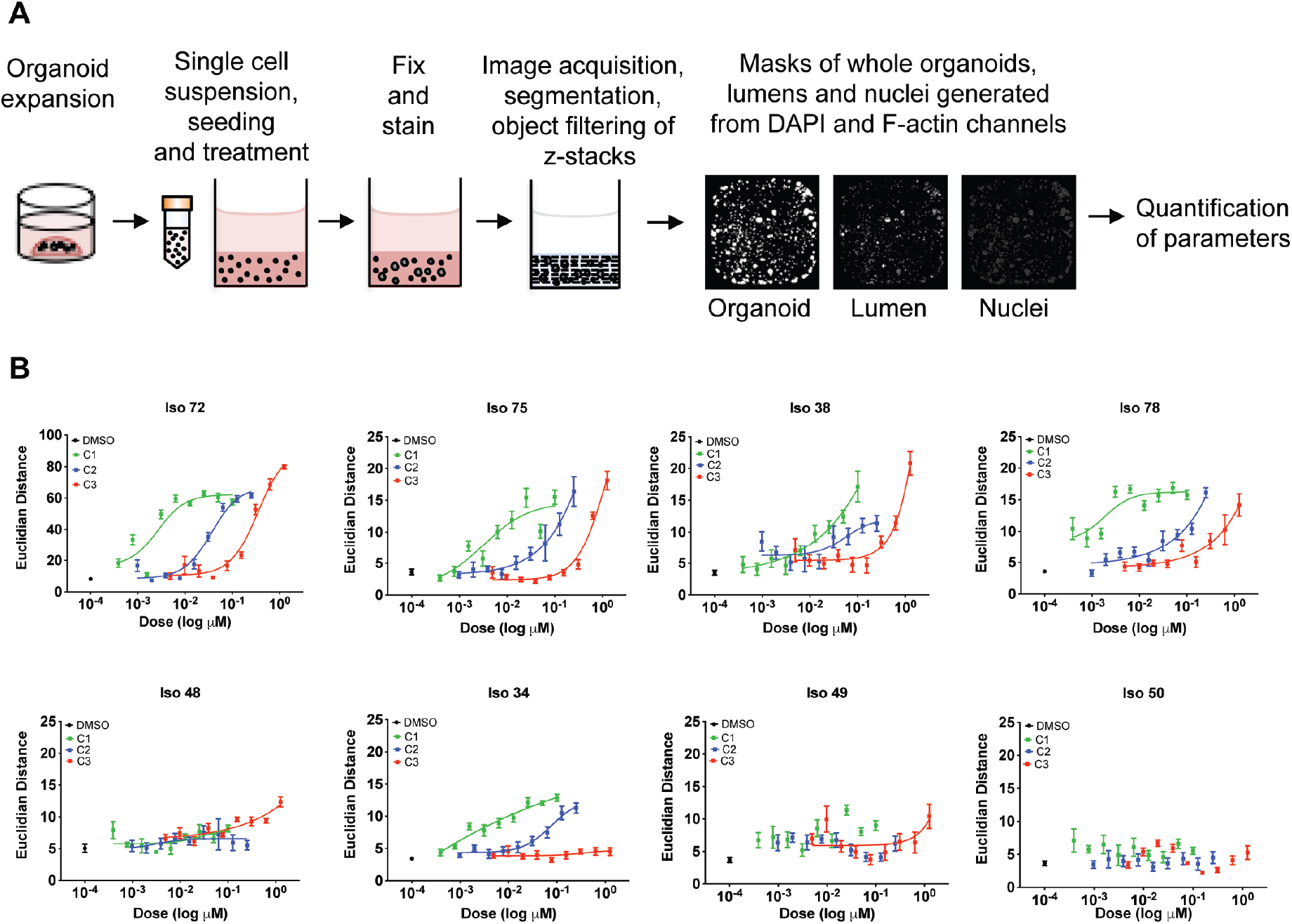
3D image analysis demonstrates target affinity of tankyrase inhibitors in organoids. **A.** Flow diagram of high-content multi-parametric image screen process. **B.** Dose response curves acquired from high content imaging screen. Organoids were seeded as single cells in 384 well plates and exposed to Tankyrase inhibitors C1, C2, C3 for six days. Individual morphological measurements were generated from DAPI and F-actin channels. Principal component analysis was used to select the top 10 most discriminating features to separate compound-induced phenotypes between high and low doses of Tankyrase inhibitors. Selected parameters were used for feature space training, whereby Euclidean distance metrics were calculated between high and low doses of each compound for each organoid line. Euclidean distances were used as an indicator of similarity between low and high doses, with a low distance indicative of high phenotypic similarity between conditions. Data are presented as mean Euclidean distance from 8 replicate wells ± S.D.

To identify optimum features that could classify TNKSi responses, feature training and Euclidian distance calculations were performed on each data set as previously described [27]. This reduced features into key differentiating components to generate a measure of distance between highest C1 TNKSi doses and negative control (DMSO, 0.1%) conditions. The feature collection that describes the compound-induced morphological change differs per organoid model, but mostly included organoid size changes, lumen shape and size, appearance of dead cells and changes in nucleus morphology related to growth reduction, cell organization and cell death induction.

The morphometric analysis showed that TNKS inhibition resulted in phenotypic alterations in Iso 38, Iso 72, Iso 75 and Iso 78 (Figure 3B). Importantly, the cellular morphometric responses correlated with known cellular EC_50_ values across a range of potencies. Compounds C1, C2 and C3 that had EC_50_ values of 2 nM, 36 nM and 319 nM also showed the expected separation of EC_50_ activities in the organoid morphometric assays, suggesting that observed effects were as a result of targeted inhibition of TNKS. Iso 72, Iso 75 and Iso 78, showed clear differential dosedependent potency for three compounds. Two additional lines, Iso 38 and Iso 34 showed partial separation of their morphological responses in line with the order of cellular potency. Iso 50, Iso 48 and Iso49 showed no discernible responses. The effect sizes of the morphometric responses compared very favourably with the noisier data from the ATP assays (Figure 2C).

Iso 75 and Iso 78 organoids required ‘full’ medium containing exogenous Wnt and R-spondin ligands for expansion in culture. Whole exome sequencing revealed that Iso 75 harboured an *RNF43* mutation (Figure 1A), which would likely yield organoids that require Wnt signalling at the receptor level – upstream of the Axin-containing β-catenin turnover complex [25]. However, both Iso 72 and Iso 38 grew independently of Wnt and R-spondin suggesting that TNKSi sensitivity is not necessarily as a result of exogenous Wnt conditions. Collectively, this data indicates that 3D image based assays provide an accurate identification and quantification of organoid responses to Wnt signalling modulators.

### Sensitivity to Tankyrase inhibitors was associated with drug-induced suppression of stem cell markers

Tankyrases promote the degradation of Axin, inducing β-catenin stabilization and activation of Wnt signalling [5]. We therefore investigated the impact of TNKS inhibition on components of the Wnt signalling pathway in the organoid lines. The cellular distribution of β-catenin was assessed by immunofluorescence in organoids treated with C1 (15 nM for 6 days). TNKS inhibition reduced total β-catenin levels in both TNKS sensitive (Iso 72 and Iso 75) and non-sensitive lines (Iso 50; Figure 4A) suggesting that TNKSi were biochemically active in all organoid lines. Quantitative RT-PCR was used to analyse gene expression changes in response to TNKSi treatment for 6 days. Expression of *DKK1* and *ASCL2*, two direct β-catenin target genes [28], were reduced in all three lines although the effect size was greater in *ASCL2* levels of the TNKSi-sensitive lines Iso 72 and Iso 75 than in the TNKSi-insensitive line Iso 50 (p<0.001, Figure 4B). Taken together, this suggested that the loss of β-catenin was linked to changes in gene expression, but not necessarily to functional responses. Changes in gene expression which did correlate with functional TNKSi sensitivity included reductions in the expression of the intestinal stem cell marker *LGR5* and the upregulation of cytokeratin 20 (*KRT20*) (Figure 4B, Supplementary figure S4), suggesting that tankyrase inhibition may induce differentiation as has previously been shown with CRC cell lines [8].

**Figure 4.**
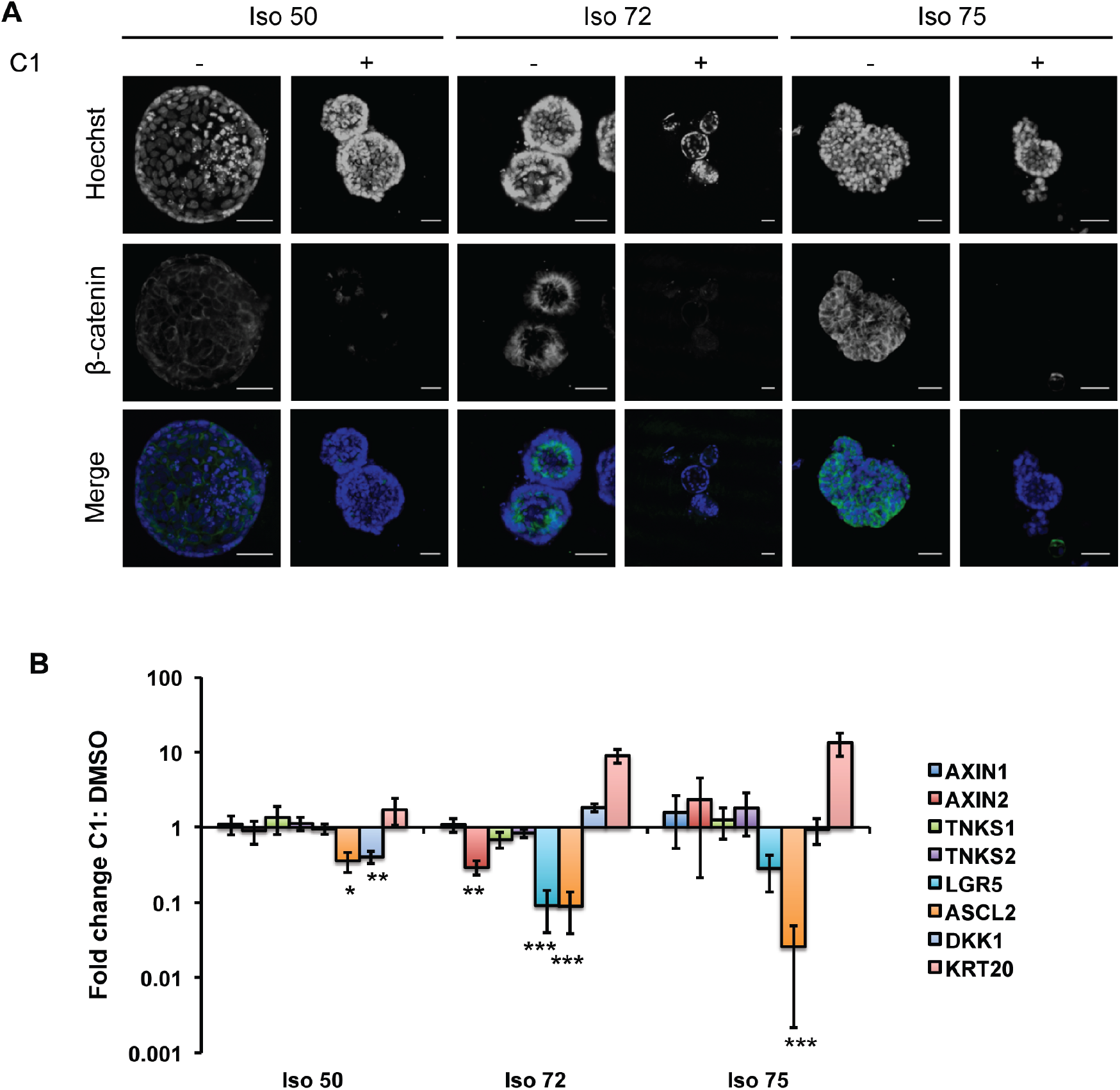
Tankyrase inhibition alters the proportion of stemness in CRC organoids. **A**. Representative confocal images of organoids stained with a β-catenin antibody following six days of exposure to C2 or control (DMSO, 0.1%) showing a reduction in the intensity and numbers of cells showing β-catenin positivity following treatment. **B**. Quantitative real-time PCR analysis of relative mRNA expression of intestinal epithelial markers from organoids (Iso 50, Iso 72, Iso 75) cultured with tankyrase inhibitor C1 (15 nM, 6 days) or DMSO (0.1%). Markers associated with intestinal stem cell activity (LGR5, ASCL2), differentiation (KRT20, DKK1) alongside other members of the Wnt–signalling pathway were determined, with mRNA expression normalized to control conditions. Experiments were performed in biological repeats (n = 3) with error bars indicative of S.E.M (***, p≤0.001; **, p≤ 0.01; *, p≤0.05, Paired Student’s t-test with Bonferroni correction.)

Lgr5 immunostaining in the Iso 50, Iso 72 and Iso 75 lines correlated with the qRT-PCR results (Supplementary Figure S4). Lgr5-positive cells were detected in both TNKSi-treated and untreated Iso 50 organoids, while Iso 72 and Iso 75 to TNKSi showed reductions in expression following TNKSi treatment. Taken together, these results suggest that the pharmacological inhibition of TNKS results in an overall reduction of Wnt/β-catenin signalling in both TNKSi-sensitive and resistant organoid lines and that phenotypic responses could not be simply predicted based on the expression of biomarkers. The best overall biomarker of organoid response may instead be functional morphometric readouts, perhaps as a consequence of differential cellular responses to the compounds being studied.

### Assessing the effects of Tankyrase inhibition in a patient organoid-derived xenograft (ODX) model

The presence of cancer stem cells is often identified by transplantation of tumour cells within immune-deficient animal models and subsequent formation of tumours [29]. To assess whether both functional and phenotypic effects of TNKSi within the sensitive organoids would translate to a reduction in tumour growth *in vivo*, an organoid-derived xenograft model was generated from the TNKSi-sensitive line, Iso 75. Organoids were digested to single cells, plated in Matrigel and immediately treated with TNKS inhibitor C1 (30 nM) or vehicle control (DMSO, 0.1%) for 24 hours. Based on previous work showing rapid responses to Wnt pathway inhibition [30], it was hypothesised that the pre-treatment of organoids would result in a transcriptional effect as a result of Wnt signalling inhibition, altering the cancer stem cell function of cells within the organoids.

Following treatment, small organoids were injected orthotopically into the flanks of 16 gamma-irradiated Non-Obese Diabetic/Severe Combined Immunodeficiency (NOD/SCID/γ) mice, at one injection site per mouse (Figure 5A). Tumours grew from the injected organoids at 100% of sites injected. However, TNKSi pre-treated organoids showed significantly delayed tumour formation by comparison with vehicle control (Figure 5B, Log Mantel-Cox test p=0.032), suggesting that the 24 hour TNKSi exposure was sufficient to reduce the capacity of transplanted organoids to form tumours.

**Figure 5.**
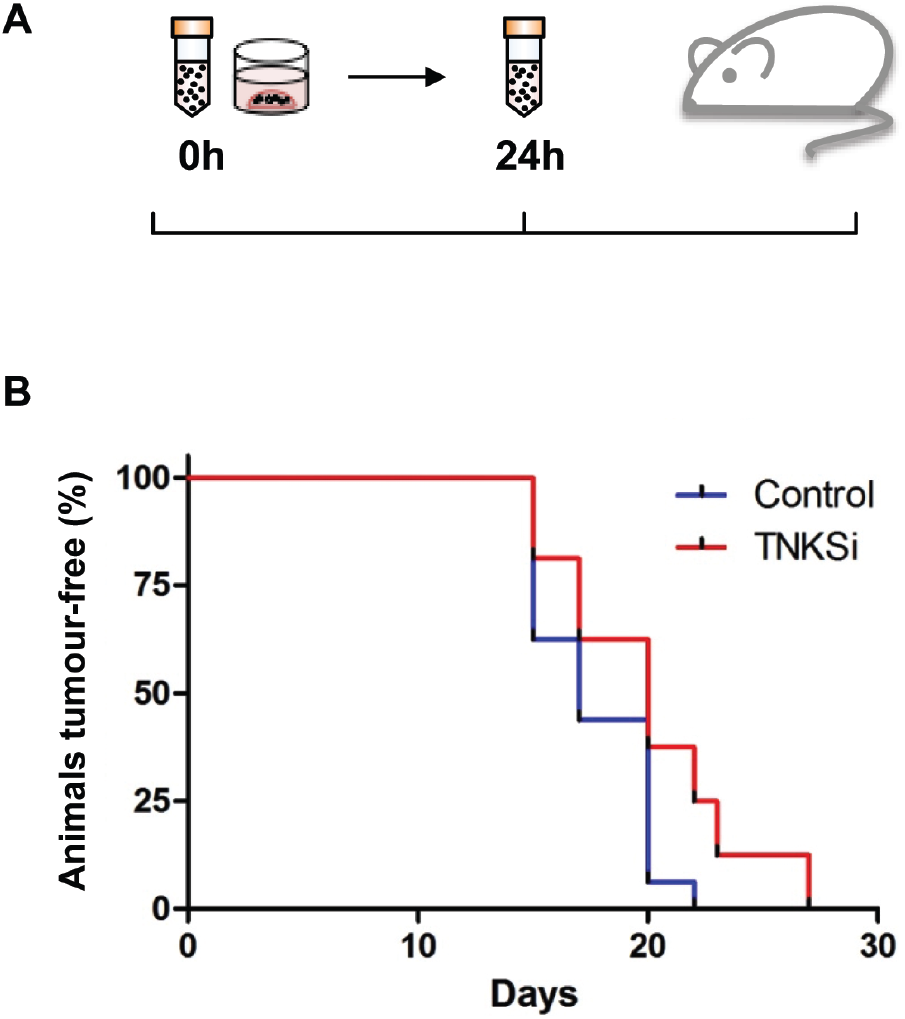
TNKSi-treated organoids enhance survival in an organoid-derived xenograft (ODX) mouse model. **A.** Schematic diagram depicting organoid transplantation into mice. **B.** A cohort of 16 NOD/SCID/ γ-irradiated mice were injected with pre-treated (30 nM C2 for 24 hours) Iso 75 organoids, or control (0.1% DMSO, 24 hours) organoids within one injection site per mouse. TNKSI-treated organoids showed a significant delayed formation of tumours compared to control conditions (Log-Rank (Mantel-Cox) test P=0.032(*)).

## Discussion

3D organoids represent an attractive platform for the capture of the effects of therapies targeting signalling pathways involved in the regulation of complex morphometric interactions that are not well represented in clonal 2D culture. Here, compounds targeting the Wnt/β-catenin pathway were studied in patient derived CRC organoids using image-based analyses as previously described [26, 27]. Numerous morphometric features, such as the distance of the matrix to central lumens and the number of nuclei per structure were quantified using automated algorithms. Due to their biological complexity, multi-parameter sets of organoid-responses may be able to characterise treatment-induced phenotypes that are linked to underlying signalling and cell biological processes. For example, changes in the activity of regulators of cell lineage will likely be reflected in alterations to the spatial organisation of cell types and organoid morphology organoids [31]. However, predictions as to how any particular morphological response is driven by alterations to specific biological mechanisms will require the development of predictive multiscale mathematical models of organoid morphology and signalling pathways [32].

Overall, morphometric analyses were found to improve assay windows by comparison with biochemical assays, most likely by aggregating many phenotypic responses that manifest themselves within a large set of morphometric parameters that can be combined into sets that sensitively discriminate responses to drug treatments using multiparametric analyses [26, 33].

The identification of a set of morphometric parameters that accurately distinguish between compounds of different potency allowed putative biomarkers of TNKSi sensitivity to be compared with organoid phenotypic responses. Single parameter functional readouts such as TNKSi-induced alterations in ATP levels showed a reduced effect size by comparison with the morphometric responses. Alterations in a range of alternative biomarkers correlated poorly with morphometric responses. Alterations in the expression of stem cell marker genes including *ASCL2* and *LGR5* correlated with cellular responses, but larger-scale studies will be required to determine whether panels of gene expression markers reliably predict organoid (and ultimately patient) responses. Confounding factors in these analyses are likely to be the levels of stem cell gene expression, variation in the ‘stemness’ of tumours and the proportion of cancer stem cells within tumour and corresponding organoid populations. Each of these properties themselves are likely to be dynamically altered following compound treatment, requiring detailed time-course analyses to deconvolute. By contrast, changes to developmental trajectories induced by compounds such as TNKSi will leave a permanent record in each organoid’s morphological structure.

Morphometric assays showed that Iso75, a line that required Wnt3a and R-spondin for growth (mutant for *RNF43* mutant and APC wild type), was sensitive to TNKSi as expected [25]. However, a simple correlation between Wnt-dependence in culture and TNKSi sensitivity was not observed since 5 out of the 8 lines examined showed morphometric responses that tracked the cellular potency of the TNKS inhibitors (Figure 3B). In particular, clear phenotypic responses were observed in Iso 72 and 78, both of which were APC mutant and only one of which required Wnt/Rspo-conditioned medium for growth.

Whilst it is mechanistically challenging to directly link Wnt inhibition via morphometric responses to alterations in stem cell cycling, pre-treatment of a TNKSi sensitive organoid line for 24h *in vitro* delayed subsequent tumour formation *in vivo* following transplantation. Taken together with the reductions in Wnt target gene expression and β-catenin levels, this would suggest that modulation of Wnt signalling in a sensitive line was sufficient to limit the stem-like signature of organoid cell populations.

Despite the benefits of using the morphometric assay formats described here to identify clear responses to Wnt modulating therapies, key challenges remain to be addressed. From a clinical perspective, further detailed investigation of Wnt modulators will be required to determine whether TNKS inhibitors can provide clinical benefit. Toxicity remains an issue with inhibitors [8], however recent studies have suggested these might be reversible [34], such that careful timing of Wnt-inhibitor treatment in combination with standard-of-care therapies could identify rational therapeutic opportunities. When strategies targeting cancer stem cells have been further developed, a robust *in vitro* readout such as organoid morphology may be able to identify those patients who would benefit the most. Lastly, the imaging pipeline described might be usefully applied in drug discovery screens, enabling the detection of compounds that might otherwise have been rejected for lack of efficacy during pre-clinical drug testing.

## Materials and Methods

### Materials

Corning Growth factor-reduced Matrigel was purchased from VWR (#734-1101). Cell culture media were purchased from Invitrogen Life Technologies, and cell culture plastics from Nunc unless otherwise stated. All TNKSi compounds were supplied and synthesized by Merck Healthcare KGaA (Darmstadt, Germany). All stock solutions of compounds were reconstituted in DMSO. Activity of each compound had been assessed prior to shipping using a cell-based immunobead assay to determine TNKSi potency based on the ability of the inhibitors to stabilise the AXIN2 [35]. The determined EC_50_ values of 2 nM, 36 nM and 319 nM (Figure 2A) were determined for C1, C2, and C3 respectively.

### Human Tissue

Surgically resected patient materials were obtained from University Hospital of Wales by the Wales Cancer Bank with informed ethical consent and anonymised (WCB project reference #12/001). The Wales Cancer Bank has ethics approval as a Research Tissue Bank from the Wales Research Ethics Committee 3 (reference 16/WA/0256), and is licensed by the Human Tissue Authority under the UK Human Tissue Act (2004) to store human tissue, taken from the living, for research (licence 12107) [36]. These approvals cover the collection of samples (including consent), processing and storing samples across multiple collection and storage sites. All patient derived material was handled in concordance with HTA regulations. Histological sections of patient tissue were imaged at University Hospital Wales and remained anonymised.

### Organoid culture

The isolation of tumour organoids from patient material was processed as previously described by Sato *et al*., [20] with some refinements outlined below. Briefly, following surgical resection, tumour samples were obtained and stored in Hibernate A medium (Life Technologies) at 4°C. Tissue pieces were dissected to remove connective tissue and chopped to pieces approximately 2 mm in diameter. Following dissection, carcinoma pieces were washed in PBS then digested enzymatically with Collagenase and Dispase for 30 minutes at 37°C. Once digested, tissue was triturated in PBS at room temperature to release cell fragments from tissue. Cell fragments within the supernatant were then centrifuged for 5 min at 100 rcf, at 4°C. Cell pellets were resuspended in Advanced DMEM/F12 supplemented with 1 % GlutaMAX, 1% HEPES buffer solution and 1% Penicillin/Streptomycin, then filtered (70 μm). Collected tissue fragments were counted and plated at a density of 1000 cell fragments per 50 μL of Growth Factor Reduced Matrigel (BD Biosciences) in 24 well plates (Nunc). Following Matrigel polymerisation, cells were overlaid with 500 μL of either “7+” or “Full” media and replenished every 4 days. ‘7+’ Media conditions consisted of Advanced DMEM/F12 supplemented with penicillin/streptomycin, 2mM GlutaMAX, 10 mM HEPES,1 X B27 supplement, 1 X N2 supplement (all Life Technologies), 1 mM N-acetyl-L-cysteine (Sigma-Aldrich). The following additional niche factors were added to generate “Full” media: 50 ng/ml mouse recombinant EGF (Life Technologies), 100 ng/ml mouse recombinant Noggin (Peprotech), 10% RSpo-1 conditioned medium [37], 40% Wnt-3A conditioned medium [20], 500 nM A83-01 (Tocris) and 10 μM SB202190 (Sigma). Y-27632 (Tocris) was also added to media for the first three days of culture. Optimal media was determined for each tumour organoid line on the basis of conditions that favoured the most efficient growth of organoids. For subsequent passaging, maintenance in culture and analyses, the most favourable media condition was used. All culture were maintained in humidified incubators at 37°C, 5% CO_2_.

### DNA extraction

DNA extraction from organoid cultures was carried out using a QIAamp DNA Mini Kit (Qiagen) following the manufacturer’s instructions. Purified DNA quality was initially assessed using a Nanodrop1000 (Thermo Fisher). Patient blood samples were processed by the Wales Cancer Bank (WCB), genomic DNA was extracted from 4 ml of whole blood using the ChemagicSTAR automated cell lysis and DNA extraction workstation (Hamilton Company) housed within the All Wales Medical Genetics Service (AWMGS) laboratory. Both DNA samples were then passed to the Wales Gene Park for quality control, library preparation and sequencing.

### Whole exome library preparation

The following was carried out by the Wales Gene Park. Genomic DNA was initially quantified using the Qubit^®^ (Life Technologies) and the samples were serially diluted to 5 ng/μL. 50ng of gDNA was used as the input and the sequencing libraries were prepared using the Illumina^®^ Nextera Rapid Capture Enrichment kit (Illumina Inc.). The steps included tagmentation of gDNA, clean-up of the tagmented DNA, amplification, clean-up of the amplified DNA, hybridisation of probes, capture of the hybridized probes, second hybridization of probes, second capture, clean-up of the captured library, amplification of enriched library, clean-up of the enriched library and finally validation of the complete library. The manufacturer’s instructions were largely followed with extra quantitation steps prior to the hybridization of the probes to ensure that approximately 500 ng of each sample was pooled. The libraries were validated using the Agilent 2100 Bioanalyser and a high-sensitivity kit (Agilent Technologies) to ascertain the insert size, and the Qubit^®^ (Life Technologies) was used to perform the fluorometric quantitation. Following validation, the libraries were normalized to 10 nM, pooled together and clustered on the cBot™2 following the manufacturer’s recommendations. Pools were then sequenced using a 75-base paired-end (2×75bp PE) dual index read format on the Illumina^®^ HiSeq4000 according to the manufacturer’s instructions.

### Whole exome sequencing and mutation calling

Reads were mapped to the human reference genome (GRCh37) with BWA version 0.7.10 [38], and duplicate reads were removed with Samtools version 0.1.19 [39]. Base quality score recalibration was performed using GATK’s BaseRecalibrator (GATK version 4.0.3.0, [40, 41] and GATK’s Mutect2 was used to call somatic variants (tools found at https://software.broadinstitute.org/gatk/). Variants were annotated with Variant Effect Predictor version 91.3 [42], using a cached version of ensemble v91. A custom Perl script was used to extract genes of interest from annotated variant files. Default parameters were used for all software unless otherwise stated. All of the calls reported were cross-checked against the cBioportal database (https://www.cbioportal.org) and are those found to have been previously described within the COSMIC database (https://cancer.sanger.ac.uk/cosmic) as being “likely oncogenic”.

### Immunohistochemistry

Formalin-fixed paraffin embedded histological sections of patient tissue were processed, imaged and analysed by the Histopathology unit at University Hospital Wales. For organoid histology, organoids embedded within Matrigel were fixed with 4% paraformaldehyde for 15 min at room temperature. Organoids were then lifted from wells and washed several times with PBS prior to embedding within low melting-point agarose, followed by dehydration, paraffin embedding, sectioning and Hematoxylin+Eosin (H&E) staining. Images were acquired on an Olympus Dp26 microscope.

### Western Blot detection

Organoids were harvested from 6 X 50 μL blobs of Growth factor-reduced Matrigel using Cell Recovery Solution (Corning) for 1 hour on ice. Total protein was extracted from organoids by addition of 500 μL of lysis buffer to whole organoids (0.02 M Tris-HCl, 2 mM EDTA, 0.5% v/v NP-40 (IPEGAL) in ddH2O containing 1x PhosSTOP phosphatase inhibitor (Roche) and 1x complete Protease Inhibitor Cocktail (Roche)). The lysates were centrifuged at 8000 rcf for 15 min and proteins harvested in the supernatants. Protein content was measured using a BCA protein assay kit (Pierce). Samples were resolved on Novex NuPAGE 4–12% Bis-Tris PAGE gels, and then blotted onto nitrocellulose membranes using the Invitrogen iBlot Dry Blotting cassette system.

Blots were blocked at room temperature for 1 h in 5% w/v milk powder in wash buffer, Tris buffered saline (TBS) plus 0.2% Tween 20. Primary antibodies were diluted in blocking buffer and incubated overnight at 4°C. After washing the blots were incubated with HRP-conjugated secondary antibodies for 1 h at room temperature. Bands were visualized using the enhanced chemoluminescence (SuperSignal West Dura; Pierce). Antibodies and dilutions: TNKS1/2 (Santa Cruz (Tnks1/2 E-10) SC-365897, 1:500), Axin2 (Cell Signalling Technologies 76G6, 1:500) (# 2151)), β-actin (Sigma (AC-74) A2228, 1:10,000), anti-mouse or antirabbit HRP-conjugated antibodies (GE Healthcare, 1:5000).

### Organoid viability measurements

Organoids in culture were gently dissociated to near-single cell populations using TrypLE (Life Technologies) before resuspension within growth factor-reduced Matrigel, and dispensed into white clear bottomed 96 well plates in 9 μL Matrigel per well (400 cells/ μL of Matrigel). Following Matrigel polymerization, a 9-point 2-fold dilution range of compounds were diluted in tailored growth medium and dispensed within each well. DMSO was used as a negative control at a final concentration of 0.1%. Following a six day exposure to compounds, Cell Titer Glo 3D (Promega) reagent was applied as per the manufacturer’s guidelines for an endpoint readout. Relative Luminescence values were obtained on a BMG Fluostar plate reader. Dose response curves and EC_50_ values were obtained using Microsoft Excel with XLFit plug-in (Version 5.4.0.8).

### High content fluorescence microscopy

Organoid lines previously established in culture were split to single cells using TrypLE (Life Technologies) and seeded within growth factor-reduced Matrigel (Corning) in black clear-bottomed 384 well plates, at 12 μL per well, at a density of 400 cells/ μL Matrigel. Upon polymerisation of Matrigel, growth media containing a titration range of individual compounds or DMSO controls were added to each well prior to incubation at 37°C. Organoids were exposed to compounds for a total of six days prior to fixation in a fix-and stain solution containing 4% para formaldehyde (Sigma Aldrich), 0.2% Triton X-100 (Sigma Aldrich) 0.25μM rhodamine-phalloidin (Sigma Aldrich) and 1 μg/ml Hoechst 33258 (Sigma Aldrich) in PBS at 4°C for 24 h. After staining, plates were washed with PBS and sealed. Imaging was performed on an ImageXpress^®^ high-content micro confocal platform (Molecular Devices). A total of 25 Z-stack images were acquired at 10 μm steps through the focal plane per well of a 384 well plate using a 4 X objective. Image stacks were captured from both TRITC channel (EX=548, EM=645) and DAPI channel (EX-380, EM=435) to detect both rhodamine-phalloidin (F-actin) and Hoechst (nuclei).

### High content image analysis

Images obtained using the ImageXpress platform were processed for phenotypic analysis within OMiner™ software (OcellO B.V.) integrated within the KNIME Analytics Platform (Konstanz, Germany, http://www.knime.org/) as described previously [26, 43]. Briefly, this software enabled the quantification of z-stack images derived from Hoechst and rhodamine-phalloidin stained organoids. Information from both the DAPI and TRITC channels were used to enhance organoid boundary separation. Masks generated from both channels facilitated the visualisation of main organoid structure, internal morphometries such as individual lumens, as well as nuclei per organoid. Objects that were out of focus were filtered and discarded from analysis.

Feature training between the negative control organoids and the high dose TNKSi treated organoids was performed to condense the phenotypic features and calculate the Euclidian distance. The results are plotted with GraphPad Prism 7 (GraphPad Software, La Jolla, CA). Results are shown as means ± standard deviations unless otherwise stated.

### Quantitative Real-time PCR

Total RNA was extracted from organoids resuspended in trizol solution (Life Technologies) containing 125 μg/ml glycogen. cDNA was yielded from purified mRNA using ImProm II Reverse Transcription kit (Promega). Quantitative-RT PCR was then performed using the SensiFAST SYBR Green Hi-ROX master mix (Bioline). Primers were designed (Sigma Aldrich) and listed in **Supplementary Table S3**. All samples were measured in triplicate, with gene expression normalised to GAPDH housekeeping gene. Reactions were measured on a Step One Plus realtime PCR instrument (Applied Biosystems) with relative changes in gene expression calculated using the ΔΔCt method.

### Wholemount immunofluorescent staining

Organoids were seeded as single cells in 96 well plates in 9 μL growth factor reduced Matrigel at a density of 400 cells/μL. Following a six day exposure to compounds and control conditions, organoids were fixed in 4% PFA for 15 min at room temperature. Wells were then washed with PBS containing 100 mM glycine prior to blocking overnight in PBS containing 10% horse serum, 0.1% Bovine Serum albumin, 0.2% Triton X-100, 0.05% Tween 20 then stained with primary antibodies overnight (Supplementary Table S3) at 4°C. Secondary antibodies were then added overnight at 4°C, prior to counterstaining with Hoechst. Wells were washed with PBS then imaged on a confocal microscope, Leica TCS SP2 AOBS.

### *In vivo* assessment of TNKS inhibitor efficacy

All mice were obtained commercially from Charles River and then housed under UK Home Office regulations. The work described here was carried out under Home Office PPL 30/3279. *In vivo* organoid engraftment studies (also termed “organoid derived xenografts” (ODX)) were conducted using immune-deficient NOD/SCID gamma irradiated mice. For organoid transplantation, organoids were in culture for 24 h from single cells prior to subcutaneous transplantation into the right flank of the animal (one injection site per mouse). Mice were checked every 2 days for tumour growth by palpation, according to Home office regulation protocol. Palpable tumours (> 5 mm) were counted for use in Kaplan-Meier analysis.

**Table 1.** Summary of EC_50_ values obtained by morphometric analysis.

| EC<sub>50</sub> (nM) | C1 | C2 | C3 |
| --- | --- | --- | --- |
| Cell line | 3 | 36 | 300 |
| Iso 34 | 3 | 75 | No Fit |
| Iso 38 | 9 | 55 | 2000 |
| Iso 48 | No fit | No fit | No fit |
| Iso 49 | No fit | No fit | No fit |
| Iso 50 | No fit | No fit | No fit |
| Iso 72 | 2.6 | 36 | 346 |
| Iso 75 | 3 | 31 | 1060 |
| Iso 78 | 1.8 | 30 | 400 |

## Abbreviations

CRC: Colorectal cancer
NOD/SCID: Non-obese Diabetic/Severe Combined Immunodeficiency
ODX: Organoid derived xenograft
PARP: poly (ADP-ribose) polymerases
TNKS: Tankyrase

## Author Contributions

Conception and design of the study: L.M.B, A.J.H, A.R.C, D.W, P.S, T.C.D, Acquisition of data (including acquisition and management of patient material.): L.M.B, A.J.H, B.H, K.B.E, J.R.S, D.A.B, C.E., H.-P.B, R.H, D.W Analysis and interpretation of data (including statistical analysis, biostatistics, computational analysis): L.M.B, A.J.H, B.H, K.Y, K.B.E, J.R.S, D.A.B, M.N, K.A, L.S.P, Drafting, review and/or revision of the manuscript: L.M.B, A.J.H, T.C.D, Study supervision: P.S, T.C.D

## Acknowledgements

We acknowledge all members of the University Hospital of Wales (UHW) Colorectal surgical team and UHW Histopathology team. The authors would like to thank the Wales Cancer Bank for providing access to all patient material and to patients for providing their consent to this study. We also acknowledge members of the Bioimaging facility at Cardiff School of Biosciences for technical support. We acknowledge our colleagues at the Wales Gene Park for their insight and expertise that assisted this research, and their technical and bioinformatic support in generating the NGS data. Wales Gene Park is a Health and Care Research Wales funded infrastructure support group. We acknowledge the guidance and expertise of Laura Thomas, Hannah West and Elena Meuser of the Inherited Tumour Syndrome Research Group within the School of Medicine, Cardiff University with bioinformatic workflows and CRC genetics. We acknowledge the support of Cellesce Ltd. We thank Victoria Marsh Durban, Anika Offergeld and Mairian Thomas and for manuscript proof reading and corrections.

