## Supplementary Figures and Data for "3D imaging of colorectal cancer organoids identifies responses to Tankyrase inhibitors"

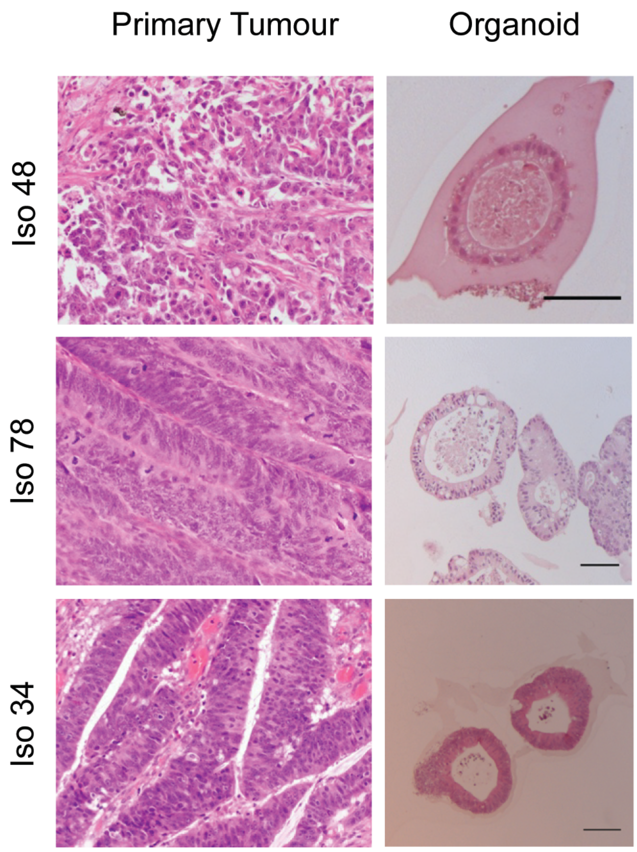


**Supplementary Figure S1**

Hematoxylin+Eosin (H & E) staining of FFPE tissue sections derived from primary colorectal tumour patient material, with organoid counterparts. Scale bar = 100 µm.

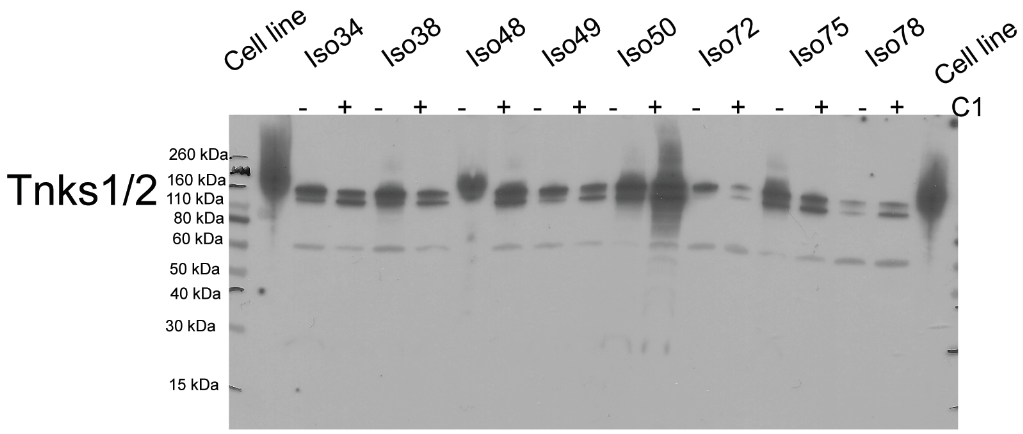


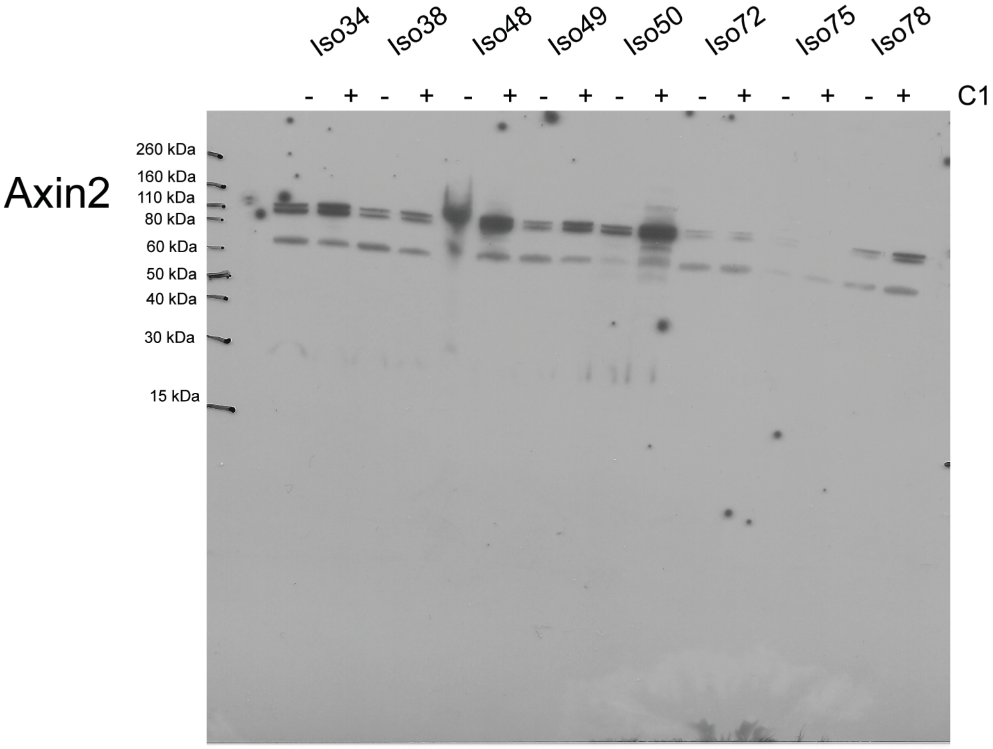


**Supplementary Figure S2**

Full scans of immunoblots. Related to Figure 2B. Organoids and cell line controls were treated with TNKSi C1 (organoids 15 nM, cell line 50nM) and blotted for TNKS1/2 top or Axin2 bottom . Novex Sharp pre-stained protein standards (Invitrogen; LC5800) were loaded as per manufacturers instructions.

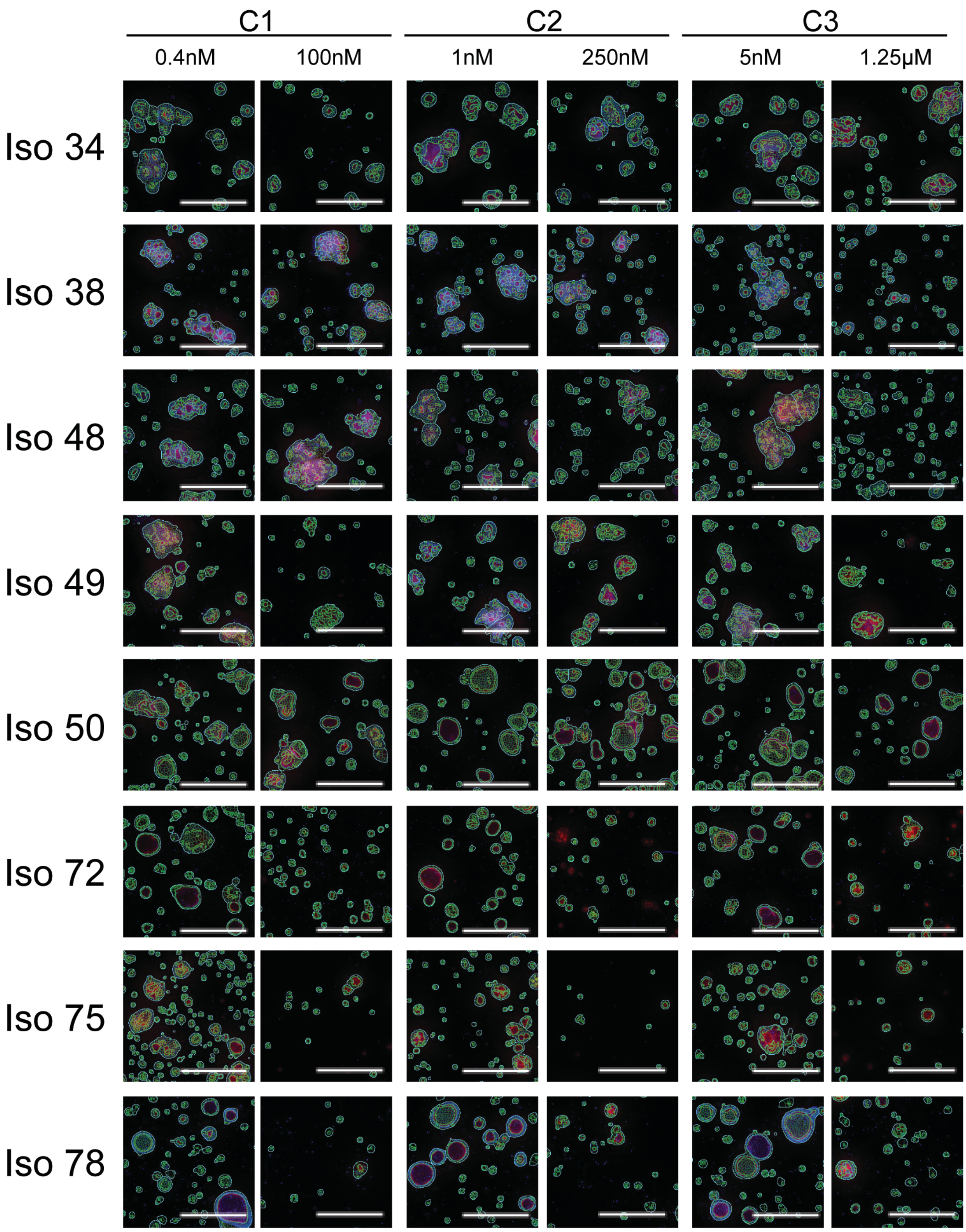


**Supplementary Figure S3**

Representative images of organoids exposed to low and high dose TNKSi (C1, C2 and C3 for 6 days) after image analysis. Projections of the Hoechst (Blue) and

Phalloidin-rhodamine (Red) signal are overlaid with the cell and lumen mask (Green). The images show 10% of the original image. Scale bar = 500 μm.

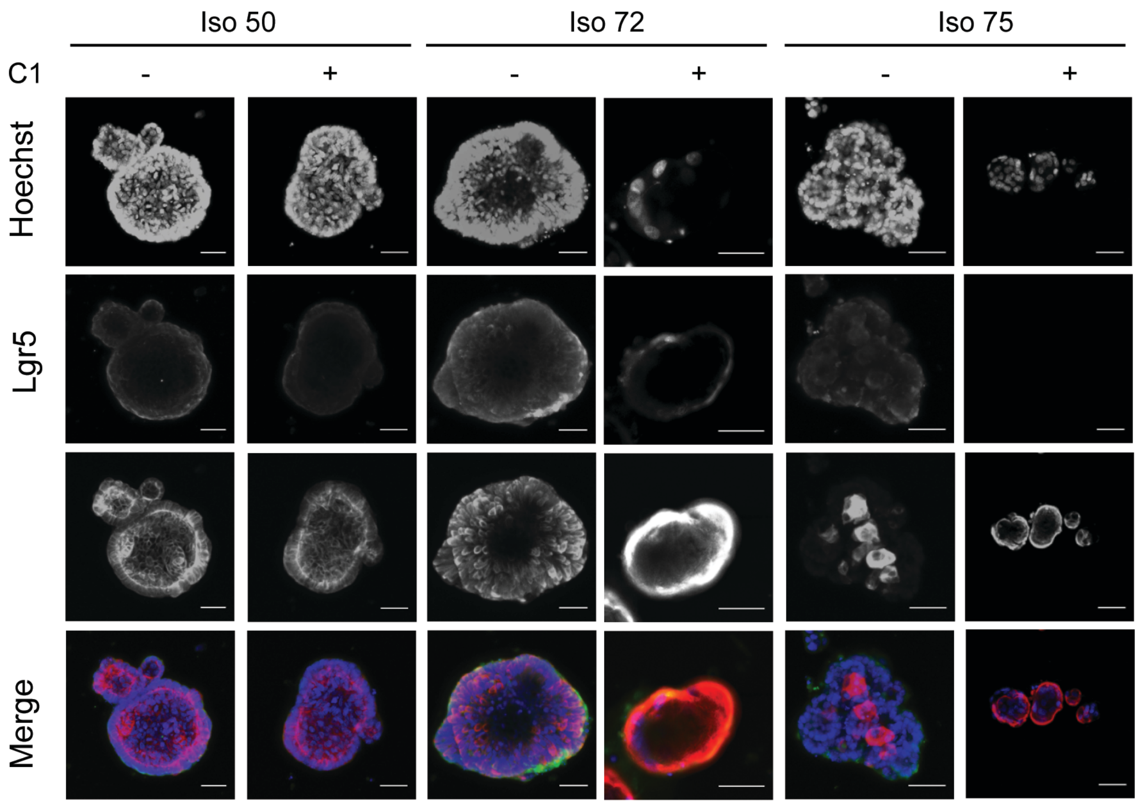


**Supplementary Figure S4**

**A**. Representative confocal images of organoids stained with an Lgr5 and Cytokeratin 20 antibody following six days of exposure to C1 (15 nM) or control (DMSO, 0.1%). Sensitive organoids demonstrated an overall reduction in the number of Lgr5 positive cells and increase in Cytokeratin 20 positive cells following treatment.

**Supplementary Table S1**

Whole Exome sequencing analysis for each organoid line. The mutations reported in the table are those found to have been previously described as likely oncogenic and curated within the COSMIC and cBioportal databases.

Supporting table are available in files Supplementary Table S1a-l

| Primary Antibody | Source | Catalogue Number | Dilution | IgG |
|---|---|---|---|---|
| Ki67 | Millipore | AB9260 | 1:100 | Rabbit |
| β-catenin | BD | 610154 | 1:250 | Mouse |
| Lgr5 | BD | 562733 | 1:250 | Rat |
| Cytokeratin 20 (KRT20) | Abcam | AB76126 | 1:100 | Rabbit |

**Supplementary Table S2.**

Primary antibodies used for fluorescence microscopy.

| Gene | Forward Sequence | Reverse Sequence |
|---|---|---|
| AXIN1 | CTGGATACCTGCCGACCTTA | CCGGCATTGACATAATAGGG |
| AXIN2 | GCGATCCTGTTAATCCTTATCAC | AATTCCATCTACACTGCTGTC |
| TNKS1 | CCGCGTGTCTGTTGTAGAGT | ACAGAAGCCCCATGCCTTAC |
| TNKS2 | TGGTGTGGGAGCCAAGTCTA | GTGGCAATTCACTCCTCTTCA |
| LGR5 | GAGTTACGTCTTGCGGGAAAC | TGGGTACGTGTCTTAGCTGATTA |
| ASCL2 | TGACCTGGGGCGTAATAAAG | ACACAGGCTTCTCCCTAGCA |
| KRT20 | ACGCCAGAACAACGAATACC | ACGACCTTGCCATCCACTAC |
| DKK1 | CCCAGGCTCTGCAGTCAGCG | CGCACGGGTACGGCTGGTAG |
| GAPDH | TGAAGGTCGGAGTCAACGGA | CCATTGATGACAAGCTTCCCG |

**Supplementary Table S3.**

Primers used for qRT-PCR by SYBR green.
