## Supplementary Table S1b for "3D imaging of colorectal cancer organoids identifies responses to Tankyrase inhibitors"

| Gene | ISO 34 | ISO 38 | ISO 48 | ISO 49 | ISO 50 | ISO 57 | ISO 68 | ISO 72 | ISO 75 | ISO 78 | Gene |
|---|---|---|---|---|---|---|---|---|---|---|---|
| APC | ▲ E1451* | ▲ G1339Ffs*2 | --- | ▲ R1450* A1446Lfs*27 | ▲ R232* E1286* | ▲ Q978* | ▲ Q1096* E1408* | ▲ Q1291* | --- | ▲ R876* E1451* | APC |
| TP53 | --- | ▲ C238Y | --- | ▲ R248Q | ▲ R248Q | ▲ R282W | ▲ R248W | --- | --- | ▲ H193D | TP53 |
| KRAS | --- | --- | --- | ▲ G12D | ▲ G12D | ▲ G13D | ▲ G13D | ▲ G12D | --- | ▲ G12D | KRAS |
| BRAF | ▲ K601E | --- | --- | --- | --- | --- | --- | --- | ▲ V600E | --- | BRAF |
| PIK3CA | --- | --- | ▲ E542K | ▲ E542K | --- | --- | --- | --- | --- | --- | PIK3CA |
| CTNNB1 | --- | --- | ▲ S33C | --- | --- | --- | --- | --- | --- | --- | CTNNB1 |
| FBXW7 | --- | --- | --- | ▲ R465C | --- | --- | --- | --- | --- | --- | FBXW7 |
| ARID1A | --- | --- | --- | --- | --- | --- | --- | --- | ▲ F2141Sfs*59 | --- | ARID1A |
| SMAD4 | --- | ▲ D537H | --- | --- | ▲ E526K | --- | ▲ Q534* | --- | --- | --- | SMAD4 |
| ARID2 | --- | --- | --- | --- | --- | --- | --- | --- | --- | --- | ARID2 |
| AXIN2 | --- | --- | --- | --- | --- | --- | --- | --- | --- | --- | AXIN2 |
| ERBB3 | --- | --- | --- | --- | --- | --- | --- | ▲ A232V | --- | --- | ERBB3 |
| MSH3 | --- | --- | --- | --- | --- | --- | --- | --- | ▲ K381Gfs*20 | --- | MSH3 |
| NRAS | --- | --- | --- | --- | --- | --- | --- | --- | --- | --- | NRAS |
| POLE | --- | --- | --- | --- | --- | --- | --- | --- | --- | --- | POLE |
| SMAD2 | --- | --- | --- | --- | ▲ S464* | --- | --- | --- | --- | --- | SMAD2 |
| TCF7L2 | --- | --- | --- | --- | --- | --- | --- | --- | --- | --- | TCF7L2 |
| RNF43 | --- | --- | --- | --- | --- | --- | --- | --- | ▲ G659GX | --- | RNF43 |
| Gene | ISO 34 | ISO 38 | ISO 48 | ISO 49 | ISO 50 | ISO 57 | ISO 68 | ISO 72 | ISO 75 | ISO 78 | Gene |
